# Revealing Complex Functional Topology Brain Network Correspondences Between Humans and Marmosets

**DOI:** 10.1101/2023.08.17.553784

**Authors:** Qiang Li, Vince D. Calhoun, Armin Iraji

## Abstract

Functional correspondences are known to exist within the brains of both human and non-human primates; however, our understanding of this phenomenon remains largely incomplete. The examination of the topological characteristics inherent in whole-brain functional connectivity bears immense promise in elucidating shared as well as distinctive patterns across different species. In this investigation, we applied topological graph analysis to brain networks and scrutinized the congruencies and disparities within the connectomes of human and marmoset monkey brains. The findings brought to light noteworthy similarities in functional connectivity patterns distributed across the entire brain, with a particular emphasis on the dorsal attention network, default mode network and visual network. Moreover, we discerned unique neural connections between humans and marmosets during both resting and task-oriented states. In essence, our study reveals a combination of shared and divergent functional brain connections underlying spontaneous and specific cognitive functions across these two species.

## I. INTRODUCTION

The configuration of functional connections in both the human and non-human primate brains has been shaped by the imperative of energy conservation and the efficient transmission of information [7, 16]. As a remarkably intricate, autonomously organizing, and remarkably energy-efficient biological neural network, the human brain exhibits distinctive patterns of information organization and connectivity when compared to its non-human primate counterparts. These two species share a fundamental reliance on two primary branches of connectivity that underlie their formidable hierarchical computational abilities: structural and functional connectivity. To delve into these properties and discern potential disparities across species, it is essential to quantify and elucidate the intricate topological attributes of the brain, an aspect that has regrettably been largely overlooked in prior research endeavors.

Our pursuit of a deeper understanding of human brain function and related brain disorders is driven by both a fundamental cognitive imperative and clinical necessity, primarily due to our high genetic affinity with non-human primate species [24]. Notably, the great apes exhibit substantial genetic similarities with humans, sharing approximately 99.9% of DNA sequences within the human population, 98.2% with chimpanzees, 97.7% with gorillas, and 96.3% with orangutans. In contrast, monkeys, encompassing macaques and marmosets, share a somewhat lower genetic overlap of approximately 93% [3].

This genetic closeness prompts a fundamental question: to what extent do these shared genetics translate into similar cognitive capabilities? Further, how do the parallels and distinctions observed in the resting and task states between humans and marmosets contribute to cognitive processes? Addressing these inquiries holds significant implications for our comprehension of intelligence across species and its relevance to advancing treatments for brain diseases [10]. In our quest to tackle these questions, we employ a multidisciplinary approach, drawing on insights from neurophysics, statistical mechanics, computational neuroimaging techniques, and the application of non-invasive functional magnetic resonance imaging (fMRI) for the measurement and mapping of brain connections [18].

Marmosets, in particular, are frequently utilized as preclinical models for studying brain diseases due to the high degree of conservation in brain-related genes between marmosets and humans [24]. Consequently, investigating the intricacies of functional brain similarities and differences between humans and marmosets holds significant interest, as it allows us to discern gaps in cognitive and consciousness functions. Our study focuses on the common marmoset monkey (Callithrix jacchus), also known as the white-tufted-ear marmoset, to address these research objectives [15, 20]. It is worth noting that all human and marmoset datasets used in this research are derived from large, publicly accessible open-source repositories, ensuring compliance with international ethical guidelines. In this study, we harnessed the established cortical networks of both humans and marmosets, conducting a comprehensive comparison of the intricate brain network topologies during resting and task states. This endeavor carries profound implications for enhancing our comprehension of higher-order cognitive capacities.

## II. THEORETICAL ANALYSIS

### Resting states

The human and marmoset resting-state connectome data from the Human Connectome Project [1], which used to map human brain connectivity [19, 22], and the Marmoset Brain Mapping Project [2], which used to map marmoset brain connectivity separately [15, 20].

The human and marmoset structural and functional connections are estimated from preprocessed diffusion MRI and fMRI, based on parcellation atlases. The functional connectivity is constructed from time courses based on a pairwise Pearson correlation, i.e., the extracted brain activity from brain regions *i* and *j*, can be denoted as *X*_*i*_ = [*x*_*i*_(1), …, *x*_*i*_(*t*)], *X*_*j*_ = [*x*_*j*_(1), …, *x*_*j*_(*t*)], then correlation between brain regions *i* and *j* is represented as 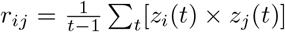, *t* is the time length of brain activity, and *z* refers to z-score value. By repeating this procedure for every pair of brain regions, the connectome matrix was generated with a size of *N* × *N*, where *N* represents the total number of brain regions. To derive structural connectivity, a series of anatomical processes are undertaken for both structural MRI (sMRI) and diffusion MRI (dMRI). Following the preprocessing and cleaning of dMRI data, alignment with anatomical data is crucial to ensuring that subsequent analyses are both anatomically informed and biologically relevant. Subsequently, the underlying white matter anatomy is mapped using diffusion tractography. With the generation of whole-brain tractography, the computation of streamline density between each region is conducted, ultimately resulting in the production of structural connectivity matrices. In summary, the construction of structural connectivity involves the following steps: 1.) Align the anatomical (T1w) image, 2.) Generate Freesurfer surfaces and parcellations, 3.) Map the human (Schaefer 200 atlas) and marmoset (MBMv3 atlas) cortical atlas to the surface, 4.) Preprocess dMRI data, 5.) Perform diffusion tractography, 6.) Generate network matrices from the regions of the human and marmoset brain atlases. Finally, we will have a *H*_*SC*_*/H*_*FC*_ = 200^2^ and *M*_*SC*_*/M*_*FC*_ = 192^2^ connection matrix for humans and marmosets, as shown in Figs. 1 2, where *H* refers to humans and *M* refers to marmosets.

**FIG. 1.**
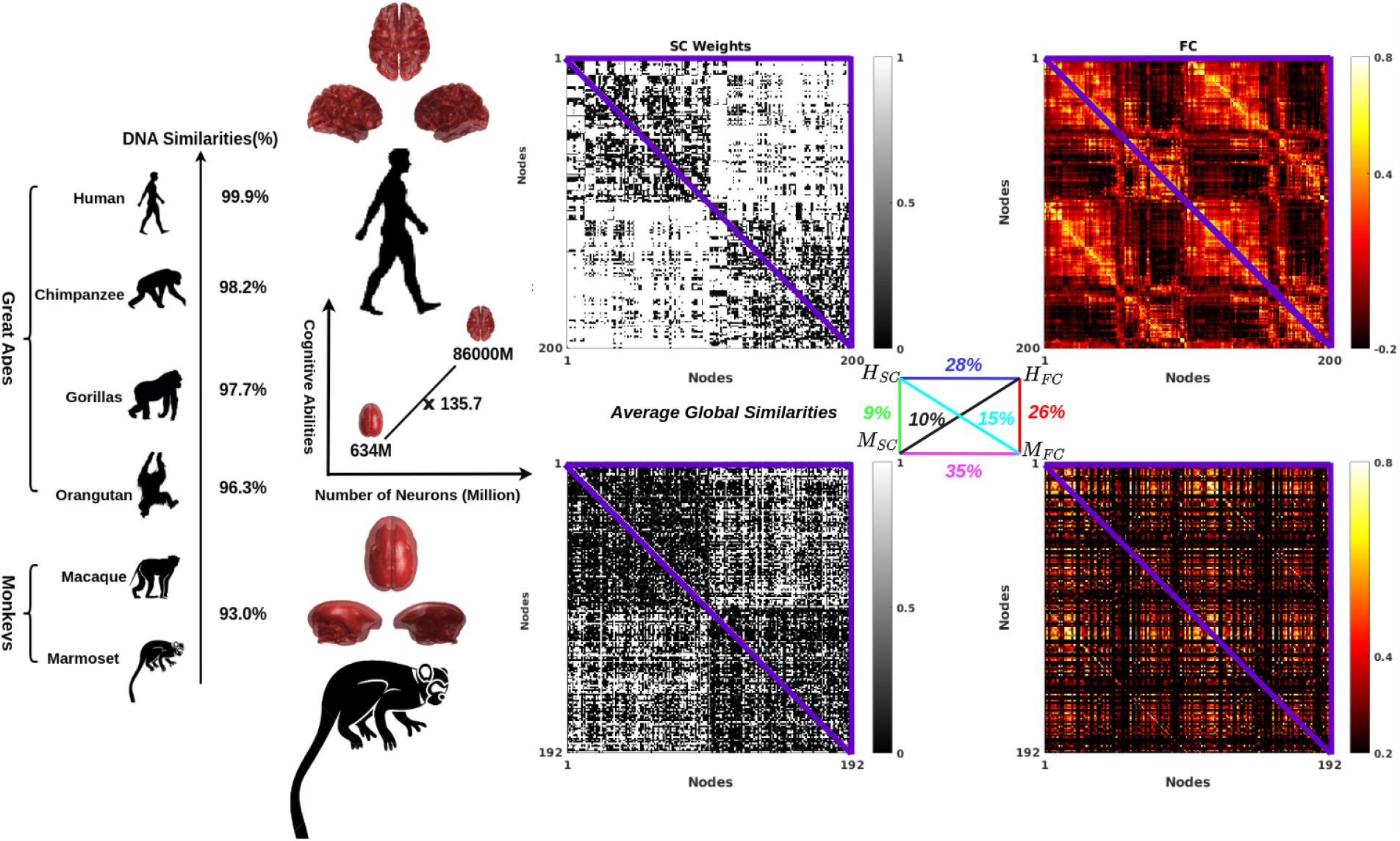
The humans (*H*) and marmosets (*M*) shared similarities and differences in genetics, i.e., DNA sequences, number of neurons, and structural (*H*_*SC*_,*M*_*SC*_) and functional (*H*_*FC*_,*M*_*FC*_) connectivity. From a deeper perspective, the great apes and monkeys have high gene overlap compared to humans, but they have large differences in their number of neurons, i.e., 634M vs. 86000M (× 135.7), and their brain structural and functional connections, i.e., 28%, 35%, 9%, 10%, 15% and 26%.

**FIG. 2.**
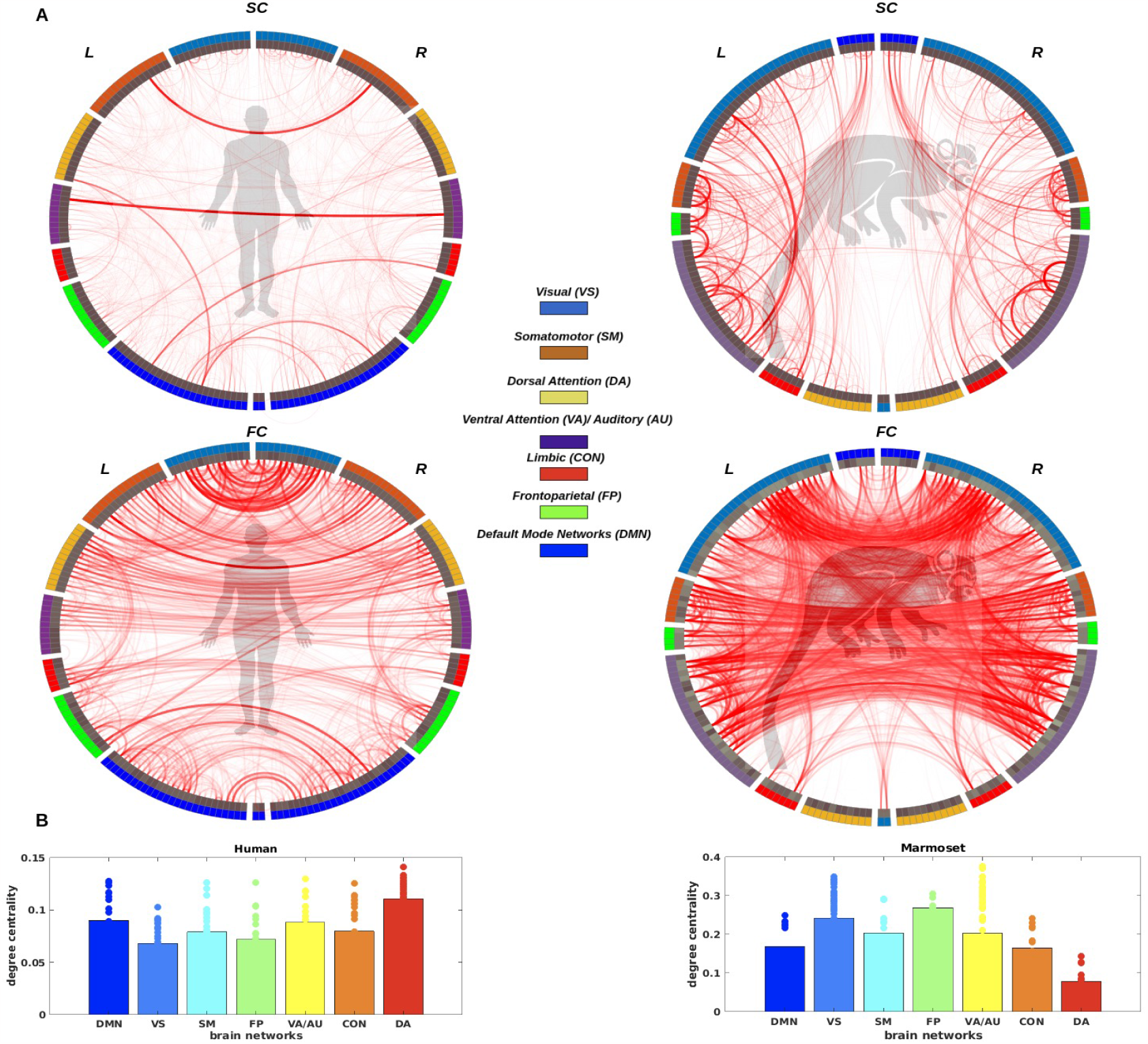
The connectogram construction and visualization scheme are identical to structural connectivity and positive functional connectivity across species, where the brain regions were categorized into seven brain networks, as shown in (A). Additionally, within the second inner circle, there was an observed manifestation of degree centrality. The related degree of centrality was quantified across brain networks in both species, as shown in (B).

As we mentioned before, physically structured connections constrain primate brain functional connectivity, and they have a moderate relationship, as shown in Fig. 1. Humans and marmosets have similar mean global Pearson correlations for both structural and functional connectivity, i.e., *ρ*(*triu*(*H*_*SC*_), *triu*(*H*_*FC*_)), 28% and 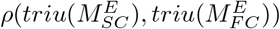, 35%, respectively. However, they share small similarities across species in structural and functional connectivity, i.e., 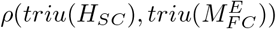, 15% and 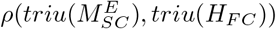, 10%, and furthermore, structural and functional connectivity also have small global similarities between two species, i.e., 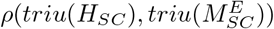, 9% and 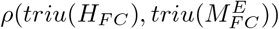, 26% separately, where *E* refers to the expend matrix of the same size as the human connectome matrix given the variance in the number of regions, an interpolation technique was applied to the marmoset structural and functional connectivity data. The mean value of marmoset structural and functional connectivity was computed, followed by the selection of a random value in close proximity to this mean, then this iterative process to obtain the final result. It is essential to note that in the initial configuration, both structural and functional connectivity encompassed comprehensive brain regions for both humans and marmosets. These regions were aligned, ensuring the inclusion of crucial regions in the connectome matrix. The result shows that humans and marmosets share large functional connectome similarities, but they still have a large gap, and it may be suggested that humans and marmosets have different spontaneous brain activity in the resting state.

To quantify disparities in the topology of brain networks across species, we applied the concept of degree centrality to both human and marmoset functional connectivity, as it furnishes valuable insights into macroscopic brain network organization [8]. Specifically, when degree centrality exhibits a high value, it signifies that certain brain regions are characterized by a greater number of connections with other brain regions, thus exhibiting a pronounced hub-like quality. Degree centrality predominantly accounts for the total number of connections associated with a given brain region [4], as illustrated in Fig. 3A.

**FIG. 3.**
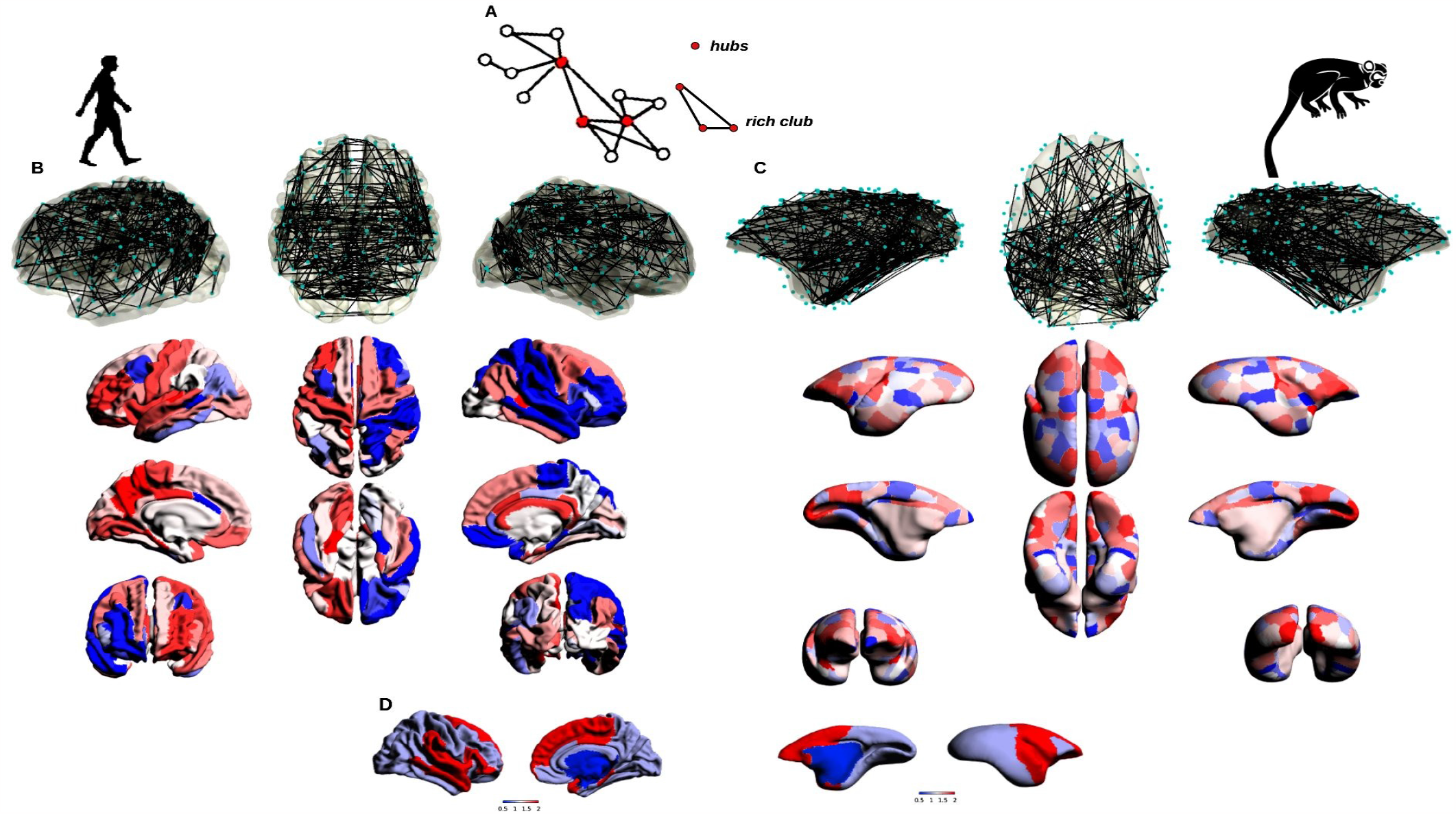
The topology of functional brain networks mapped to human primate and non-human primate brain surfaces from functional connectivity matrices is shown in Fig. 1. The hubness of human primate and non-human primate brain networks (A) (i.e., the left panel (B) is human brain networks and the right panel (C) is marmoset brain networks) is measured from above topology functional networks, and more red regions indicate stronger connectivity (rich hubness) with other brain regions, and the other way around. The network clustering with clustering number *k* = 2 was applied to both humans and marmosets, as shown in (D).

The computation of degree centrality for cortical hubs in humans and marmosets entailed the utilization of unthresholded weighted functional connectivity matrices to pinpoint regions with significant functional hub properties, as depicted in Fig.3B and Fig.3C (i.e., regions boasting a plethora of connections, and its connections correspond to Fig.2A). This was achieved by summing all pairwise functional connectivity values pertaining to each region, expressed mathematically as *C*_*D*_ = ∑_*j*_ *r*_*ij*_. As evidenced in Fig. 2B, Fig.3B, Fig.3C, and Fig. 3D, we discern certain similarities in the distribution of cortical hubs during resting states between humans and marmosets, notably within dorsal attention, default mode and visual networks. Concurrently, we may also identify variations in cortical hubs within other brain regions.

Recent methodological advancements have enabled the mapping of intricate brain properties as gradients onto low-dimensional representations [17]. These gradients have served to delineate the topographical organization of the functional brain connectome, spanning from unimodal to transmodal networks. The process involved the use of functional connection matrices to compute an affinity matrix, which captures the inter-area similarity of a specific feature. Subsequently, this affinity matrix was leveraged to unveil a gradual ordering of the input matrix within a lower-dimensional manifold space, employing dimensionality reduction techniques [23]. As visualized in Fig. 4, marmosets manifest a spectrum pattern of gradients concerning functional connections, thereby preserving the topographical configuration of functional connections akin to humans. However, discernible distinctions are apparent in the distribution of gradient spectra when comparing marmosets to humans.

**FIG. 4.**
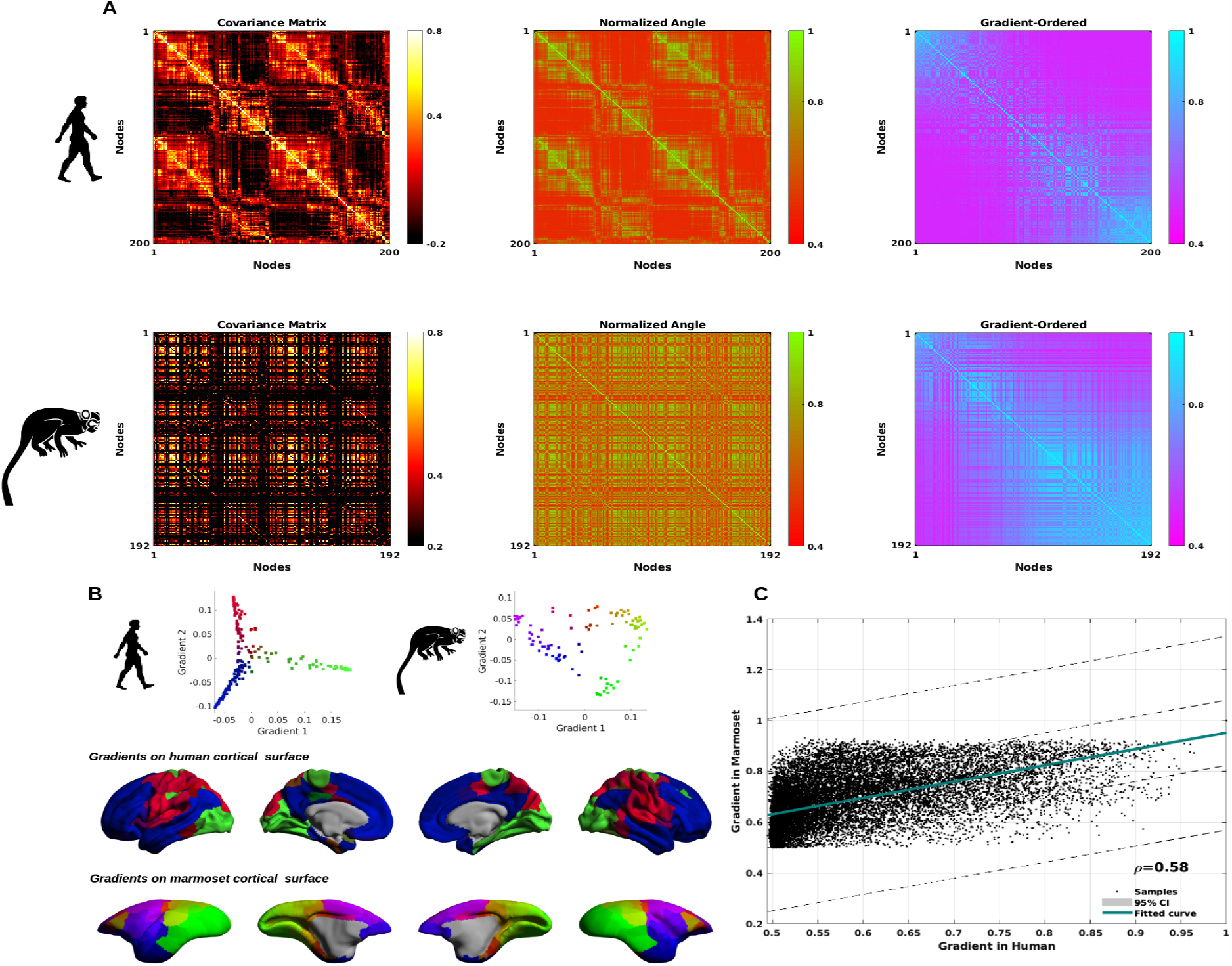
To generate gradient-based brain distribution maps, the initial step involves measuring a normalized angle map derived from pairwise functional connectivity. Subsequently, diffusion embedding [6] is applied to obtain the gradient-ordered map, illustrated in A. As we mentioned above, the functional connection matrices were utilized to generate an affinity matrix, which was then decomposed into a collection of primary eigenvectors describing the axes of greatest variance using linear rotations. The scatter plot displays the scores of each seed on the first two axes (i.e., gradient 1 and gradient 2), with colors representing position in this 2D space, i.e., green refers to visual cortex, red refers to somatosensory and auditory cortex, blue refers to association cortex, and other colors refer to the transfer from a unimodal to a multimodal state. These hues (i.e., gradient 1) can be projected back to the Conte69 hemisphere surfaces of humans [21] and the MBMv4 hemisphere surfaces of marmosets [15] individually, as shown in B. Finally, the gradient similarities across species are discussed, revealing a 58% similarity, as depicted in C.

Consequently, the parallels and disparities between humans and marmosets find expression in the gradient-based cortical topology network organization. Hence, a gradient is observed, spanning from the more sensory domains of the cerebral cortex to the more associative regions, in both humans and marmosets. To articulate this gradient disparity, one might characterize the more sensory extremity as composed of unimodal areas in marmosets, whereas the more associative extremity comprises multimodal areas.

### Task states

Exploring functional correspondences between marmosets and humans in the context of task and resting states can be achieved by examining the “fingerprints” of functional connectivity within brain regions. Resting state patterns are characterized as state-agnostic and spontaneous, making them an intriguing avenue for investigating functional correspondence. To this end, we utilized movie-driven fMRI, a methodology that allows us to directly compare the activation patterns of visual areas in marmosets and humans and establish functional correspondences within the visual cortex across both species [9].

Our analysis employed functional movie-driven task fMRI datasets, and we performed an analysis focusing on the alignment of human visual regions (FFC, FST, LO1, LO2, LO3, MST, MT, PHA, PH, PIT, POS1, PeEc, ProS, TE1p, TE2p, V1, V2, V3A, V3B, V3CD, V3, V4, V4T, V8, VMV, VVC) with marmoset-specific visual regions (A19DI, A19M, A35, A36, Ent, FST, MST, ProSt, TE1, TE2, TE3, TFO, TF, TH, TLO, TL, TPO, V1, V2, V3A, V3, V4T, V4, V5, V6) under the influence of naturalistic movie stimuli. The chosen visual regions exclusively capture responses to movie stimuli, particularly those related to face, body, and scene patches in both human and marmoset visual domains. 13 healthy volunteers, comprising 9 males and 4 females aged between 22 and 56 years, along with 8 common marmosets-six males and two females, aged between 20 and 42 months and weighing between 300 and 436 grams-participated in the study. Neural activity from human and marmoset subjects was recorded using fMRI data sourced from [9]. Subsequently, to elucidate the similarities and differences between humans and marmosets, we applied cross-correlation to the fMRI signals.

The similarity matrix was computed based on Pearson correlation, following the same approach used for resting fMRI. As depicted in Fig. 5, this similarity matrix encompasses 26 human brain regions and 25 marmoset brain regions. Notably, the maximum co-occurrence brain activity correlation observed between species reaches 52%, while the mean global co-occurrence brain activity similarity stands at 0.16/16%. Although this value may seem modest, it is essential to consider that the median similarity is 0.17, with a range spanning from -0.27 to 0.52 and a confidence interval of [0.06, 0.26]. In comparison to the resting state, our findings indicate an increase in brain activity during the task state, with the heightened functional coupling between species predominantly attributed to the visual areas of the brain.

**FIG. 5.**
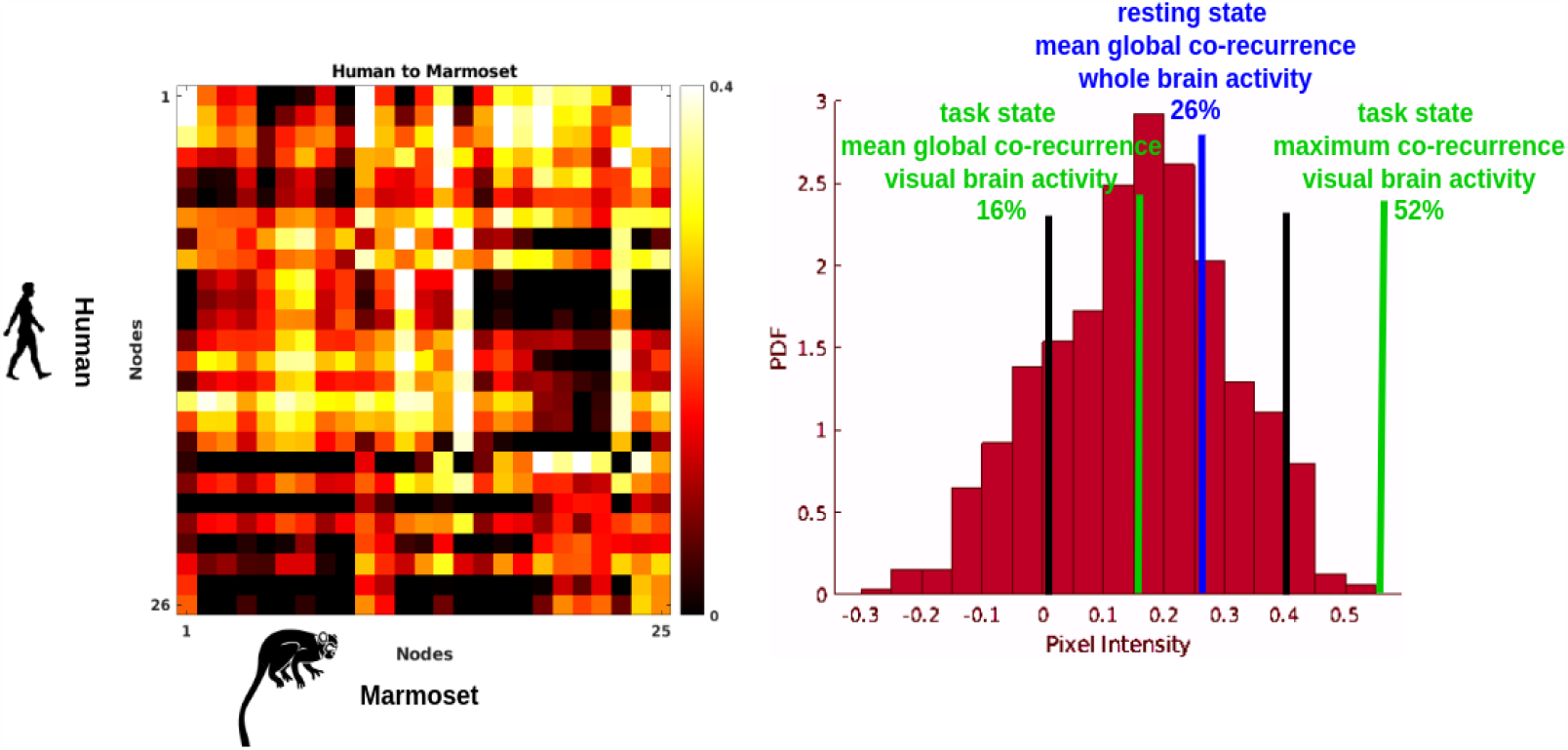
The functional connectivity similarity matrix between humans and marmosets in response to visual stimuli. In the left panel, lower-order functional connectivity based on pairwise Pearson correlation is presented for 26 human brain areas and 25 marmoset brain regions. The right panel depicts the equivalent probability density distribution (PDF). The two black lines represent the correlation matrix thresholds, while the blue line represents the mean global co-occurrence brain activity similarity between humans and marmosets in resting states (26%) and the two green lines represent the mean global co-occurrence visual brain activity similarity (16%), and maximum co-occurrence visual brain activity similarity (52%) between humans and marmosets in task states.

In summary, our investigation demonstrates a striking similarity in the arrangement of brain networks in both marmoset monkeys and humans, highlighting the conservation of higher-order visual areas across these species. While visual processing appears to be a shared trait between humans and marmoset monkeys, our findings do unveil subtle yet potentially significant functional variations. The analysis of mean global functional coupling alterations from the resting state to the task state indicates that humans and marmosets exhibit similarities in the organization of functional topology, albeit to a limited extent. However, it is important to note that the majority of structural and functional couplings differ between the two species.

These findings underscore the importance of exercising caution when employing marmosets as a computational model for comparing high-level cognitive functions in humans, despite the substantial genetic overlap and the similarities in structural and functional topology within brain functions shared by both species.

## III. DISCUSSION

This letter elucidates the investigation of both similarities and differences in brain functions between humans and marmoset monkeys, which have become widely utilized models for studying mechanisms underlying brain diseases. While humans and marmosets share substantial overlaps in their genetic sequences, rendering them a promising choice for modeling human organisms and related diseases, it is imperative to assess whether such genetic similarities extend to the functional aspects of brain connectomes. Throughout this letter, we address this crucial question by conducting a comparative analysis of complex brain structural and functional connectivity patterns in both species, employing rigorous brain network statistical analysis.

Our findings reveal the existence of functional similarities in brain functions between humans and marmosets, albeit at a moderate level. Notably, we observe pronounced disparities in functional brain networks during both resting and task states. However, it is important to acknowledge several potential limitations in our study. Firstly, the moderate global similarities observed in cross-species structural and functional coupling are derived from a direct comparison of connectivity matrices, which provides a rough estimate of correlation across species and may introduce some degree of deviation. Consequently, cautious interpretation is warranted.

Secondly, our analysis relies on estimating connections solely through pairwise Pearson correlation, which inherently overlooks higher-order information organization within the intricate brain networks. Recognizing that brain networks encompass interactions that go beyond pairwise relationships [5, 10–14], future extensions of this work should consider more sophisticated approaches to unveil potentially more intricate results.

Lastly, our statistical exploration of complex brain network topology primarily leverages graph network analysis, specifically degree centrality and cortical gradient measures. While these approaches have enabled the identification of meaningful similarities and differences across species, a more comprehensive investigation could incorporate additional quantitative metrics to fully probe the properties of information propagation within brain networks during both resting and task states.

In conclusion, our study sheds light on the multifaceted landscape of brain function comparisons between humans and marmosets, offering insights into the extent of their functional correspondence while also highlighting areas for potential refinement and expansion in future research endeavors.

## IV. SUMMARIZING

The research presented in this work reveals intricate correspondences within the complex structural and functional topology of brain networks between humans and marmosets. It is evident that structural and functional couplings within each species display substantial similarities, but when considering cross-species comparisons, both structural and functional connectomes exhibit no-table distinctions. This divergence is noticeable not only during resting states but also in task-cognitive states.

The application of network science techniques has allowed us to quantify these complex brain networks and has unveiled the presence of rich hubness and gradient distribution, particularly pronounced in dorsal attention, default mode and visual networks, within both species. In summary, our findings underscore the existence of moderate brain function similarities across species, even though there is a substantial genetic overlap between humans and marmosets. Therefore, it is imperative to exercise caution when employing marmoset monkeys as a model for the study and comparison of cognitive functions with humans.

This work was supported by NSF grant 2112455, and NIH grants R01MH123610 and R01MH119251.

